# RETINA: Reconstruction-based Pre-Trained Enhanced TransUNet for Electron Microscopy Segmentation on the CEM500K Dataset

**DOI:** 10.1101/2025.01.08.632030

**Authors:** Cheng Xing, Ronald Xie, Gary D. Bader

## Abstract

Electron microscopy (EM) has revolutionized our understanding of cellular structures at the nanoscale. Accurate image segmentation is required for analyzing EM images. While manual segmentation is reliable, it is labor-intensive, incentivizing the development of automated segmentation methods. Although deep learning-based segmentation has shown promise, its accuracy to automatically segment cellular structures in EM data remains insufficient compared to expert manual results. Current approaches usually use either convolutional or transformer-based neural networks for image feature extraction. We developed the RETINA method, which combines pre-training on the large, unlabeled CEM500K EM image dataset with a hybrid neural-network model architecture that integrates both local (convolutional layer) and global (transformer layer) image processing to learn from manual image annotations. RETINA outperformed existing models on cellular structure segmentation on five public EM datasets. This improvement works toward automated cellular structure segmentation for the EM community. Data and code are available at [Zenodo](https://zenodo.org/records/12795779?preview=1&token=eyJhbGciOiJIUzUxMiJ9.eyJpZCI6IjRkOGI1OTc4LTk1MjUtNDQ5ZC1iYTcxLWI0MjRmODMxMjI0NyIsImRhdGEiOnt9LCJyYW5kb20iOiJmMGI0MDE4NTNlNTFkY2Q2ZTRmYzM2ZDIxM2EwYmQxMSJ9.hy10jz1RrEYE9qKCVntHGXb1lI9n30uRoEi-Fg1QMeafT6d8L3cLXRpdRkGw8qrrtwyu87lqQFJ5eWLJg-3aVA) and [GitHub](https://github.com/BaderLab/RETINA).

## 1 INTRODUCTION

Electron microscopy (EM) has revolutionized our understanding of cellular structures at the nanoscale (Mü ller et al., 2024). Three-dimensional volume EM enables detailed visualization of cells, cellular structures and entire organisms (Peddie et al., 2022; Hoffman et al., 2020; Vergara et al., 2021; Witvliet et al., 2021). Image segmentation (marking objects and regions of interest) and annotation (labeling objects and regions) is required for EM image analysis and interpretation. Manual segmentation of EM datasets is reliable but labor-intensive, incentivizing the development of automated EM image segmentation methods (Aswath et al., 2023). Deep learning-based segmentation is a promising solution for automated image segmentation (Ma et al., 2024b), but its accuracy on subcellular structures in EM data is not sufficient to replace manual segmentation by a human expert (Gallusser et al., 2022).

Supervised deep learning approaches have traditionally been used for image segmentation, where manual identification of pixels that are part of a given type of object are used to train a model to predict the segmentation of unseen pixels. This is limited by the availability of manual training data. An improved approach is to use transfer learning to transfer model parameters from similar problems, not necessarily based on EM images. For example, the ImageNet dataset of millions of images (Deng et al., 2009) can be used to pre-train a model and transfer general image processing ability to new data and tasks (Minaee et al., 2021). Although ImageNet pre-trained models provide a solid starting point, domain differences in their training data may impact their ability to extract fine-grained features effectively in nanoscale images (Shrestha et al., 2023; He et al., 2019). Research suggests that creating more domain-specific datasets for pre-training can substantially improve performance (Caron et al., 2019).

Fortunately, open access EM data are increasingly available in central repositories such as EMPIAR (Iudin et al., 2016) and OpenOrganelle (Xu et al., 2021). However, most EM-relevant datasets are unlabeled, limiting the use of supervised approaches. As a result, unsupervised pre-training methods, which do not require labels, have been developed and have shown promising results in transfer learning for image analysis, including for EM images (Conrad and Narayan, 2021; Chen et al., 2020a; He et al., 2020). For instance, contrastive learning enables the model to associate similar samples and differentiate dissimilar ones by predicting pairs that can be either positive or negative (Noroozi and Favaro, 2016; Chen et al., 2020a). Another effective method to improve image segmentation accuracy is with deep learning model architecture innovations, such as convolutional neural networks (CNN) (LeCun et al., 1995) and transformer architectures (Vaswani et al., 2017; Dosovitskiy et al., 2020). CNN-based models, such as U-Net (Ronneberger et al., 2015) and SegNet (Badrinarayanan et al., 2017) excel at extracting local image features (Zeiler and Fergus, 2014; Luo et al., 2016), enabled by hierarchical image feature extraction, shared weights and local receptive fields. Transformers treat an image as a sequence of patches, efficiently capturing global features (Dosovitskiy et al., 2020). Architectures that combine CNN and transformer structures to use both local and global image features can enhance segmentation accuracy. For example, SwinUNETR uses an encoder comprising Swin transformer blocks combined with a U-Net-like structure capturing hierarchical representation structure, where deeper layers in the encoder correspond to decreased spatial resolution and increased embedding space dimension (Hatamizadeh et al., 2021).

To improve subcellular structure segmentation in EM images, we have developed a method that combines the advantages of unsupervised pre-training on large EM-relevant datasets with a hybrid model architecture incorporating both CNN and transformer layers within the encoder. We used the CEM500K dataset, containing 0.5 x 10^6^ information-rich and heterogeneous EM images (Conrad and Narayan, 2021), for pre-training, and the TransUNet deep learning architecture (Chen et al., 2021), which features an encoder with integrated CNN and transformer layers, as our model backbone. Given the unlabeled nature of the EM images, a reconstruction-based architecture was established for TransUNet pre-training. We benchmarked our method against non-pre-trained deep learning models and previously published models pre-trained on the CEM500K dataset using MoCoV2 (Chen et al., 2020b). All models were evaluated on five public datasets: CREMI Synaptic Clefts (CREMI, 2016), Guay (Guay et al., 2021) (human platelets), Kasthuri++ (Kasthuri et al., 2015) (neocortex volume), Perez (Perez et al., 2014) (mammalian brain), and UroCell (Mekuč et al., 2020) (urothelium tissue). Our Reconstruction-based prE-trained enhanced TransUNet for electron mIcroscopy segmentatioN on the CEM500K dAtaset (RETINA) model outperforms benchmark models across these diverse datasets.

## 2 METHODS

### 2.1 Model architecture

#### 2.1.1 RETINA: pre-training

To enhance segmentation accuracy on electron microscopy (EM) data, we designed the RETINA model to pre-train the encoder on the unlabeled CEM500K dataset (Conrad and Narayan, 2021) and fine-tune it on benchmark datasets using transferred pre-trained parameters. Our model is based on the TransUNet architecture (Figure 1a), leveraging its hybrid encoder to extract both global and local features (Chen et al., 2021). We change the TransUNet architecture by replacing its segmentation head with a convolutional decoder layer that outputs a reconstructed image. This architecture also ensures compatibility with the fine-tuning phase, enabling seamless transfer of pre-trained parameters due to the matching encoder structure. Given the unlabeled nature of the CEM500K dataset, we tasked the model with reconstructing altered images to recover their original forms. This approach aids the network in learning structural details and global context by challenging it to learn the differences between the original and altered images.

**Figure 1.**
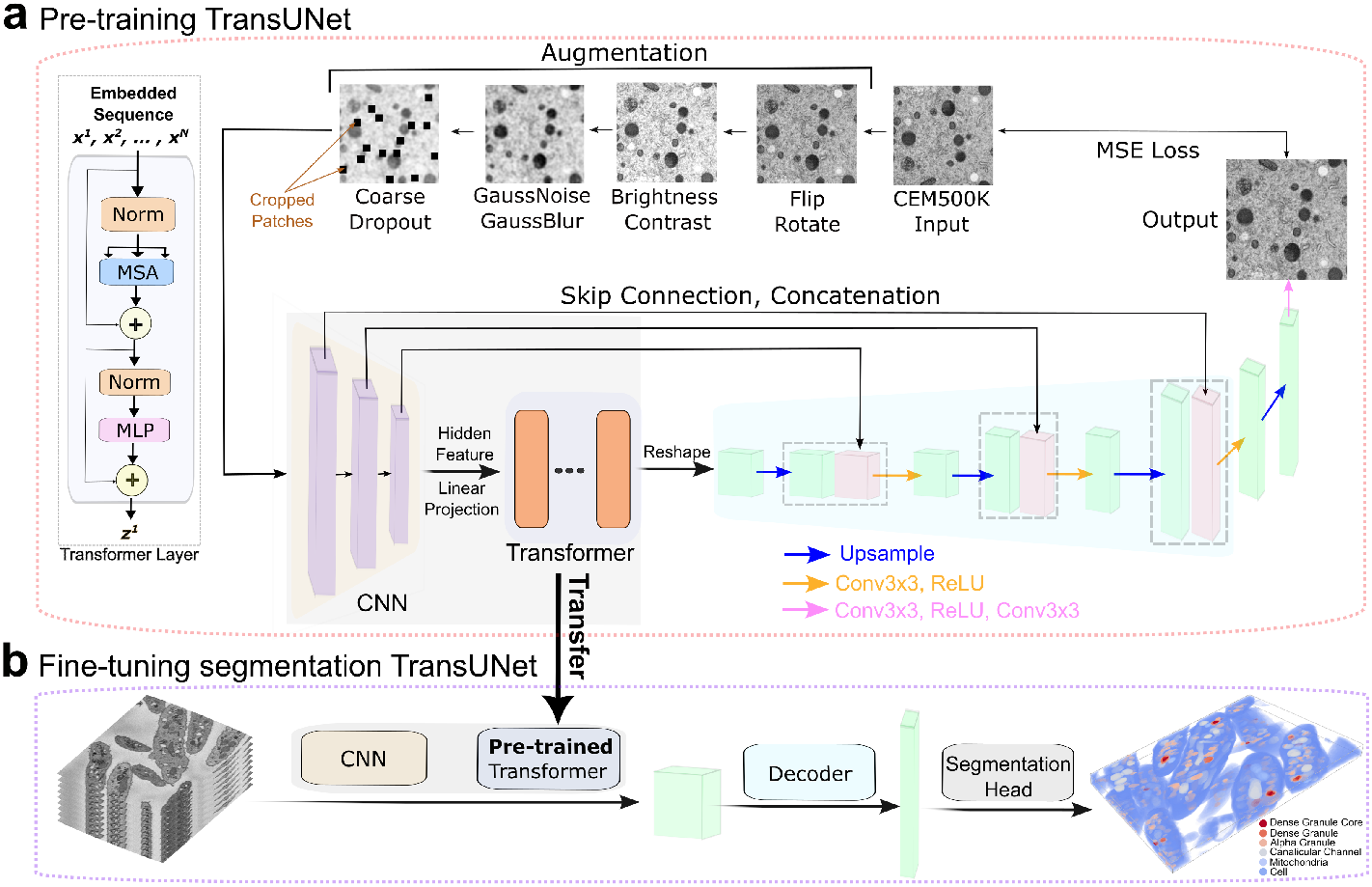
Overview of the RETINA method. **a**, RETINA pre-training process: input images from CEM500K are first augmented and then passed sequentially through convolutional and transformer layers. The embedded features are reshaped and processed by the decoder network, generating reconstructed images. The loss is computed between the input and output images. The structure of the transformer layer is highlighted on the left side. The pre-training process follows this order: augmentation, convolutional layers, transformer layers, reshaping of features, decoding, and loss computation. **b**, RETINA fine-tuning process: after transferring the pre-trained layers, 2D images are encoded by convolutional and transformer layers. The corresponding embedded features are then decoded and labeled via the segmentation head. Abbreviations: CNN, convolutional neural network; Conv, convolution; MLP, multi-layer perceptron; MSA, multi-head self-attention; MSE, mean squared error; Norm, normalization.

Initially, input images undergo a series of alterations, including flipping, rotation, brightness and contrast adjustments, noise addition, blurring, and patch cropping (Table S1) (Figure 2a). These augmentations enhance the variability of EM images and, more importantly, increase the difficulty of image reconstruction so that the model can more comprehensively pre-train the parameters of each layer. The images are then processed through the encoder, which combines convolutional and transformer layers. The convolutional layers extract high-resolution feature maps that capture local information. These feature maps are transformed into patch embeddings via trainable linear projections. The spatial information encoded in these patches is then fed into transformer layers, where multihead self-attention and multi-layer perceptron blocks enhance feature extraction capabilities (Figure 1a). Each image patch is represented by an output vector from the transformer layer, encapsulating the image’s characteristics.

**Figure 2.**
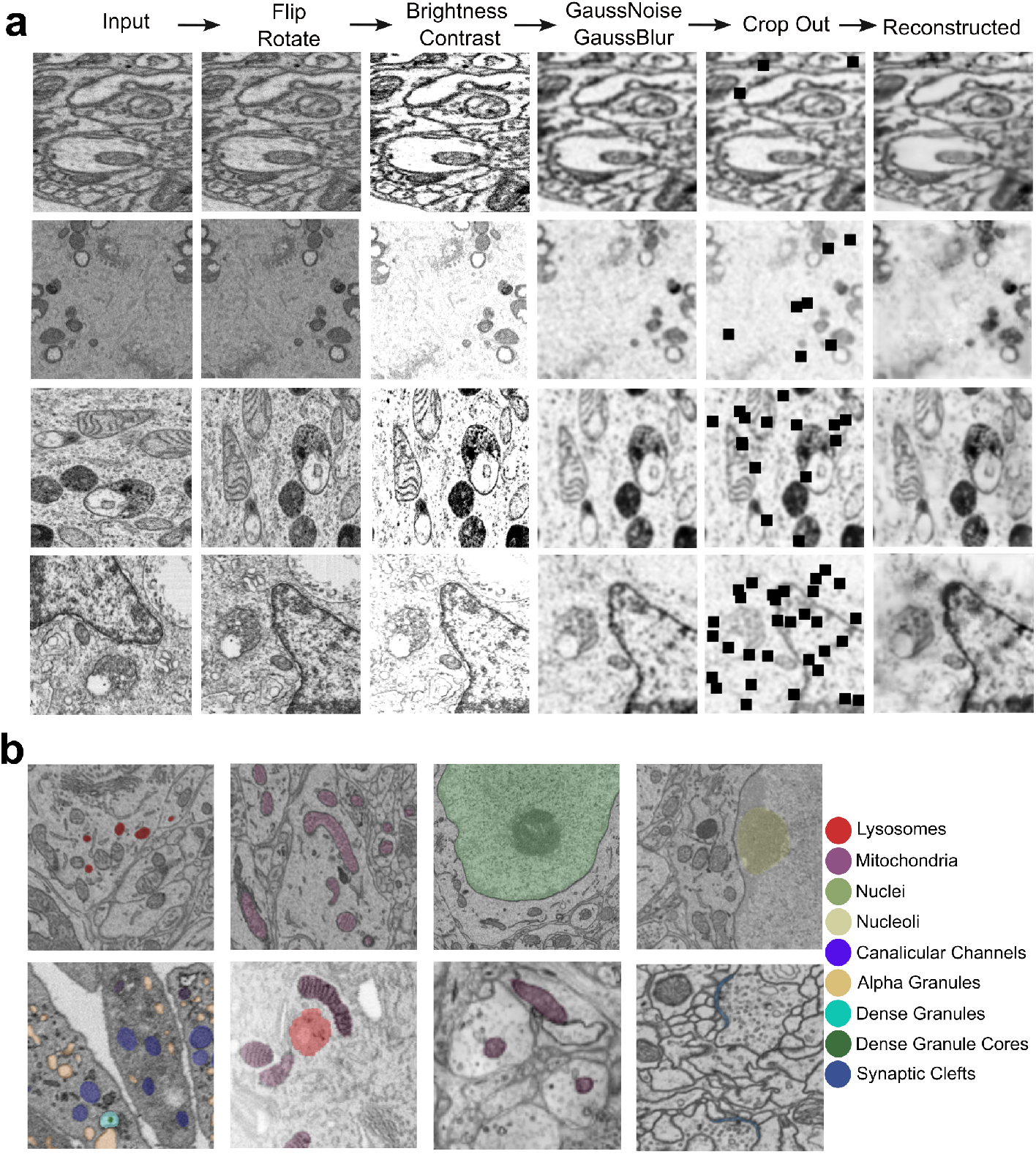
Augmented CEM500K images and benchmark datasets. **a**, Example images of CEM500K and their augmented counterparts for each augmentation step: from left to right, the columns show the Input, flip and rotate, brightness and contrast adjustment, Gaussian noise and Gaussian blur, cropping, and final reconstructed images. Each row represents an example image undergoing the entire augmentation process. **b**, Example images and label maps of each of the five benchmark datasets: the first row represents lysosomes, mitochondria, nuclei, and nucleoli in the Perez dataset. The second row corresponds to the Guay, UroCell, Kathuri++, and CREMI Synaptic Clefts datasets.

For the reconstruction task, the encoder’s extracted features are converted into a visual form by the decoder. Transitioning from the encoder to the decoder involves reshaping the matrix of patch feature vectors to recover spatial order. Subsequently, the channel size of the reshaped matrix is reduced, and the feature map is upsampled layer by layer. Skip connections, similar to those in the UNet model, facilitate the aggregation of features from different resolution levels. Finally, the feature matrix passes through convolutional layers to produce the reconstructed image. We evaluate the model output against the whole original image using mean squared error after applying same flip and rotation operation applied to the original image. The overall RETINA implementation parameters for pre-training are shown in Table S2.

#### 2.1.2 Fine-tuning using TransUNet

After pre-training, the trained parameters of the transformer layers are transferred to the corresponding transformer layers in a second TransUNet model with identical architecture to be used for fine-tuning (Figure 1b). To ensure stability during training on benchmark tasks, these transferred parameters are frozen (not updated further). However, since the fine-tuning task involves segmentation and requires pixel mask generation, the final layer is a segmentation head before the output. This adaptation of TransUNet, from reconstruction to segmentation, ensures that the model accurately performs the segmentation task while leveraging the pre-trained parameters. The overall RETINA implementation parameters for fine-tuning are listed in Table S3. Details about the model implementation can be found in the supplementary material “Implementation” section.

### 2.2 Dataset

#### 2.2.1 CEM500K used for pre-training

To improve the accuracy of EM image segmentation, we used the CEM500K dataset for model pre-training. It is an information-rich and non-redundant 25 gigabyte 2D EM image dataset (Conrad and Narayan, 2021). Briefly, 5.3 × 10^6^ images were collected and cropped into 224 × 224 pixel patches, forming the CEMraw dataset. This dataset contains a wide range of modalities and sample preparation protocols (a complete list of the datasets with related attribution can be found in the supplementary materials of the CEM500K paper (Conrad and Narayan, 2021)). The images underwent deduplication and filtering, resulting in the CEMdedup and final CEM500K datasets. The selected 0.5 × 10^6^ images that we use for training contain diverse cellular images from more than 100 unrelated biological projects (Conrad and Narayan, 2021), as shown by representative images (Figure 2a, left column).

All 0.5 × 10^6^ images in the CEM500K dataset were used as input for RETINA pre-training and processed with augmentation procedures including flipping, rotation, brightness and contrast adjustment, noise addition, blurring, and patch cropping. Flipping and rotation were randomly applied with the probability 0.5. In addition, each operation has a set of parameters (Table S1) to define the degree of augmentation. Each image went through different degrees of augmentation controlled by these parameters. This approach increases randomization to enhance data variability. For example, as shown in (Figure 2a), the first image is neither flipped nor rotated; the second is only flipped; the third is only rotated; and the fourth is both flipped and rotated. Since noise and blur were added after brightness adjustment, the images appear less bright in the fourth column. As shown in the fifth column, the number of cropped patches is randomly applied in the range from 1 to 32. For better visualization, the four representative CEM500K images input to the pre-training process are sorted according to the number of cropped patches, from the smallest to the largest (four rows in Figure 2a).

#### 2.2.2 Five benchmark datasets used for fine-tuning

We evaluated RETINA’s performance using five benchmark datasets: CREMI Synaptic Clefts (CREMI, 2016), Guay (Guay et al., 2021) (human platelets), Kasthuri++ (Kasthuri et al., 2015) (neocortex volume), Perez (Perez et al., 2014) (mammalian brain), and UroCell (Mekuč et al., 2020) (urothelium tissue) (Table S4). These datasets include nine subcellular structures for segmentation: lysosomes, mitochondria, nuclei, nucleoli, canalicular channels, alpha granules, dense granules, dense granule cores, and synaptic clefts. Representative images and ground truth labels for each dataset are presented in Figure 2b, with labels overlaid on the corresponding raw images. Guay and UroCell have multiple organelles labeled for segmentation. To consistently benchmark with previous research (Conrad and Narayan, 2021), four types of labels in the Perez dataset were split into four independent binary segmentation tasks. And the CREMI Synaptic Clefts and Kathuri++ datasets each focus on a single type of label. Additionally, these datasets involve different types of electron microscope imaging technologies (Table S4), which increases the diversity of benchmarking data.

## 3 RESULTS

### 3.1 Evaluation of RETINA on benchmark data

#### 3.1.1 RETINA achieves the best segmentation accuracy on EM benchmarks

To evaluate the segmentation capability of RETINA, we benchmarked it against several advanced models for EM image segmentation. The primary benchmark model is the UNet-ResNet50 model (He et al., 2016) pre-trained with MoCoV2 (He et al., 2020), selected for its superior performance on the CEM500K dataset compared to a randomly initialized model and an ImageNet-supervised pre-trained model (Conrad and Narayan, 2021). RETINA was subjected to 200 pre-training epochs, ensuring parity in the number of epochs with the MoCoV2 pre-trained benchmark (Conrad and Narayan, 2021). Additionally, to demonstrate the advantages of RETINA’s hybrid encoder architecture, a randomly initialized TransUNet model was included as a benchmark. Our baseline is a randomly initialized UNet-ResNet50 model, enabling us to assess the enhancement from RETINA pre-training by comparing the baseline with the MoCoV2 pre-trained model, the improvement from the RETINA encoder structure by comparing the baseline with the TransUNet model, and the integrated improvements of RETINA compared to all other models. The number of training iterations (each iteration representing a batch-level update with a corresponding learning rate adjustment) was set to ensure sufficient fine-tuning. This allowed the models to converge effectively and achieve stable Intersection over Union (IoU) scores (predicted segmentation vs. ground truth segmentation at the pixel level). We optimized the training process to minimize changes in IoU, which optimizes model performance. For consistency and comparison purposes, we selected the iteration counts based on previous work (Conrad and Narayan, 2021). The corresponding training iteration numbers are listed in Table 1. We reimplemented each benchmark model using the same code and parameters as published (Conrad and Narayan, 2021). Although the Perez dataset includes several labeled organelles, each was trained independently as a binary task and averaged to obtain the final IoU value, consistent with prior work (Conrad and Narayan, 2021). Table 1 lists the IoU values of all benchmarks and Figure S1 shows percent differences improvement of the models compared with the randomly initialized UNet-ResNet50 in IoU scores (CREMI Synaptic Clefts benchmark uses TransUNet as baseline since the IoU score was 0.000 for the randomly initialized UNet-ResNet50 model.). A high IoU score indicates better overlap of predicted segmentation results with the ground truth. The UroCell benchmark demonstrates the most significant IoU increase, over 150%, attributed to RETINA. This is mainly due to the TransUNet structure, as the TransUNet benchmark achieved nearly 150% enhancement without pre-training. However, the randomly initialized TransUNet performed over 25% worse than the baseline on the Guay dataset, but with pre-training, RETINA improved the IoU by almost 75%, indicating the benefit of pre-training. Interestingly, for the CREMI Synaptic Clefts dataset, the pre-trained benchmark model performed almost 25% worse than the non-pre-trained TransUNet, highlighting the feature extraction advantage of the hybrid model structure. For the Perez and Kasthuri++ benchmarks, RETINA still outperformed the other models, though the improvements were less pronounced likely due to the relative simplicity of these binary tasks compared to the others. Overall, the IoU scores of the fully trained models show that RETINA has the best segmentation accuracy among the benchmark models.

**Table 1.** Comparison of segmentation IoU values for RETINA versus benchmark models. The numbers of training iterations are referenced from the CEM500K publication. Lysosomes, mitochondria, nuclei, and nucleoli within Perez benchmark are listed separately. Mean values of three independent runs are reported.

| Benchmark | Training Iterations | Random Init. | Random. Init. TransUNet | CEM500K moco | RETINA |
| --- | --- | --- | --- | --- | --- |
| CREMI S.C. | 5000 | 0.000 | 0.294 | 0.246 | **0.327** |
| Guay | 1000 | 0.313 | 0.390 | 0.467 | **0.511** |
| Kasthuri++ | 10000 | 0.904 | 0.894 | 0.915 | **0.916** |
| Perez | 2500 | 0.850 | 0.893 | 0.905 | **0.919** |
| Lysosomes | — | 0.845 | 0.845 | 0.854 | **0.885** |
| Mitochondria | — | 0.841 | 0.847 | 0.887 | **0.890** |
| Nuclei | — | 0.984 | 0.990 | 0.990 | **0.991** |
| Nucleoli | — | 0.731 | 0.891 | 0.889 | **0.910** |
| UroCell | 1000 | 0.203 | 0.579 | 0.598 | **0.610** |

#### 3.1.2 RETINA converges quickly

In many application areas, models can be found to achieve strong performance with a few fine-tuning steps if they are pre-trained (Minaee et al., 2021). We tested if this holds with RETINA by evaluating the performance (IoU) changes of all tested models and datasets at different numbers of fine-tuning iterations (Figure 3a). In terms of method comparison using pre-training with different model architectures, we evaluated RETINA against the MoCoV2 pre-trained model across five benchmark datasets. On the CREMI Synaptic Clefts dataset, RETINA achieves an IoU of 0.330 within 1000 iterations, maintaining optimal performance with minimal additional gains in subsequent iterations (Figure 3a). In contrast, the MoCoV2 pre-trained model reaches comparable results by 3000 iterations. On the Kathuri++ dataset, both pre-trained models stabilize by 400 iterations, but RETINA reaches a higher IoU value earlier. For the simpler binary segmentation of the Perez dataset, RETINA reaches an optimal state early on, while the MoCoV2 pre-trained model converges rapidly but lags 100 iterations behind RETINA (Figure 3a). On the UroCell dataset, RETINA attains an IoU of over 0.6 within 200 iterations, whereas the MoCoV2 pre-trained model is slower to converge, improving from 400 to 800 iterations (Figure 3a). Overall, RETINA converges more rapidly than the MoCoV2 pre-trained model in most experiments.

**Figure 3.**
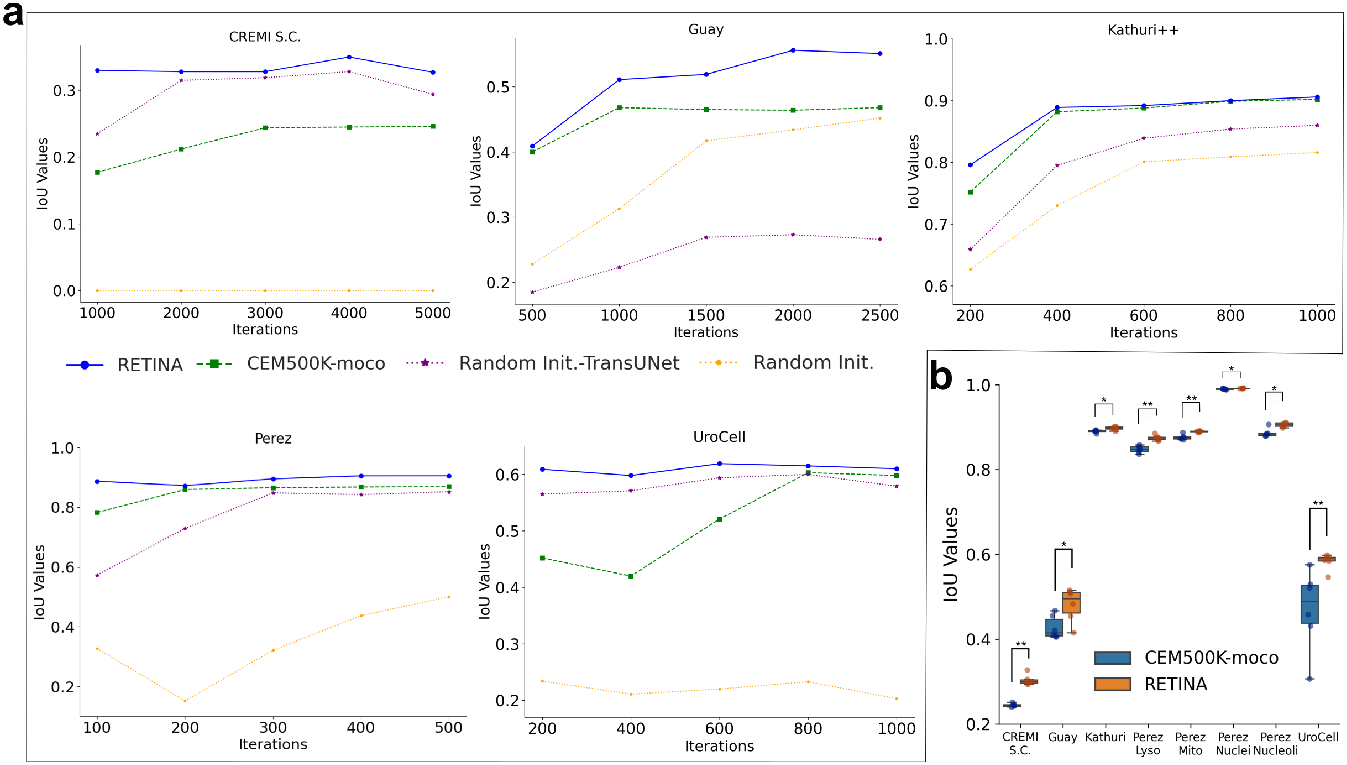
RETINA demonstrates fast convergence and robustness, outperforming other models across all segmentation benchmarks. **a**, IoU values of RETINA and other benchmark models across different fine-tuning iterations. Each IoU value is obtained by inferring on test datasets using model weights trained with the specified number of iterations. **b**, Box plots representing the distribution of IoU values for each benchmark, with each benchmark consisting of six experiments corresponding to different random seeds. Models pre-trained with MoCoV2 on CEM500K are colored in blue, and RETINA is in orange. *p<0.05, **p<0.01.

To assess RETINA against non-pre-trained benchmark models, we compared it to randomly initialized TransUNet and UNet-ResNet50 across five datasets. On the CREMI Synaptic Clefts dataset, the non-pre-trained TransUNet requires 2000 iterations to reach peak performance, 1000 iterations slower than RETINA. The randomly initialized UNet-ResNet50 baseline consistently produces an IoU of 0.000 (Figure 3a), as previously reported (Conrad and Narayan, 2021), underscoring the difficulty of this dataset and rendering it unsuitable for convergence speed comparisons. On the Guay dataset, RETINA converges around 2000 iterations, whereas the non-pre-trained UNet-ResNet50 model exhibits slower convergence up to 2500 iterations. TransUNet stabilizes at approximately 1500 iterations, but with much lower accuracy, making convergence comparisons less informative (Figure 3a). For the Kathuri++ dataset, non-pre-trained models continue to improve up to 600 iterations or beyond, while RETINA reaches stability earlier. On the Perez dataset, both non-pre-trained models take longer to converge compared to RETINA. On the UroCell dataset, the non-pre-trained TransUNet continues gradual improvement up to 600 iterations (Figure 3a), though the convergence slope is low. The UNet-ResNet50 baseline performs poorly throughout the 200–1000 iteration range, making meaningful comparisons on this dataset less relevant. Across most experiments, RETINA converges faster than both random initialization methods. In summary, RETINA achieves optimal performance more quickly across five benchmark datasets compared to all other models tested.

#### 3.1.3 RETINA is robust to random seed changes

RETINA exhibits superior segmentation accuracy on EM datasets by leveraging pre-training on CEM500K and using the TransUNet backbone. The initial experiments, however, were conducted using a single random seed across all models. To assess the reproducibility of our results, we repeated the fine-tuning of RETINA and the MoCoV2 pre-trained benchmark model on each dataset using six different random seeds. The resulting IoU score distributions demonstrate that RETINA consistently significantly outperforms the MoCoV2 pre-trained model across all datasets (Mann-Whitney U test, Figure 3b). Notably, RETINA exhibits substantially lower variance in the UroCell multiclass benchmark, underscoring its ability to reduce uncertainty in more complex multi-class segmentation tasks. Across the other benchmarks, RETINA maintains consistently low variance across different random seeds. Overall, RETINA significantly outperforms the MoCoV2 pre-trained model in accuracy and is robust to random seed variation.

#### 3.1.4 RETINA evaluation by manual image inspection

To visually assess segmentation quality, we manually inspected representative label maps for each benchmark. We reviewed images in two and three dimensions (2D and 3D). For 2D viewing, we took the images as used in the segmentation task. For 3D viewing, we selected the Guay and CREMI Synaptic Clefts datasets, as they contain representative nanoscale labeled structures: Guay includes seven different organelles, and the CREMI Synaptic Clefts dataset contains a unique label not found in other datasets. All visualizations were generated from the models used in the RETINA benchmarking experiments detailed in Table 1.

For the 2D inferred visualization, we manually identified improved segmentation regions among the models, demonstrating that RETINA produces output masks closest to the ground truth in the first row (Figure 4a, dashed-outline squares). For example, comparing with the ground truth images (top row), RETINA successfully identifies lysosomes highlighted in cyan in the last row of Figure 4a; however, the TransUNet (second row) and MoCoV2 pre-trained model (third row) are not able to detect the organelles in these highlighted regions. While nucleoli segmentation in the Perez dataset shows no significant differences among the models, RETINA excels in the relatively challenging regions of the Kathuri++ dataset, where both TransUNet and the MoCoV2 pre-trained model fail (yellow squares). RETINA also accurately identifies the lysosome region marked with blue squares, whereas TransUnet identifies less area and the MoCoV2 pre-trained model misassigns the region as mitochondria (purple) instead of lysosomes (red). RETINA’s ability to more thoroughly identify entire target objects and avoid mislabeling helps explain its higher IoU values compared to other benchmark models (Table 1).

**Figure 4.**
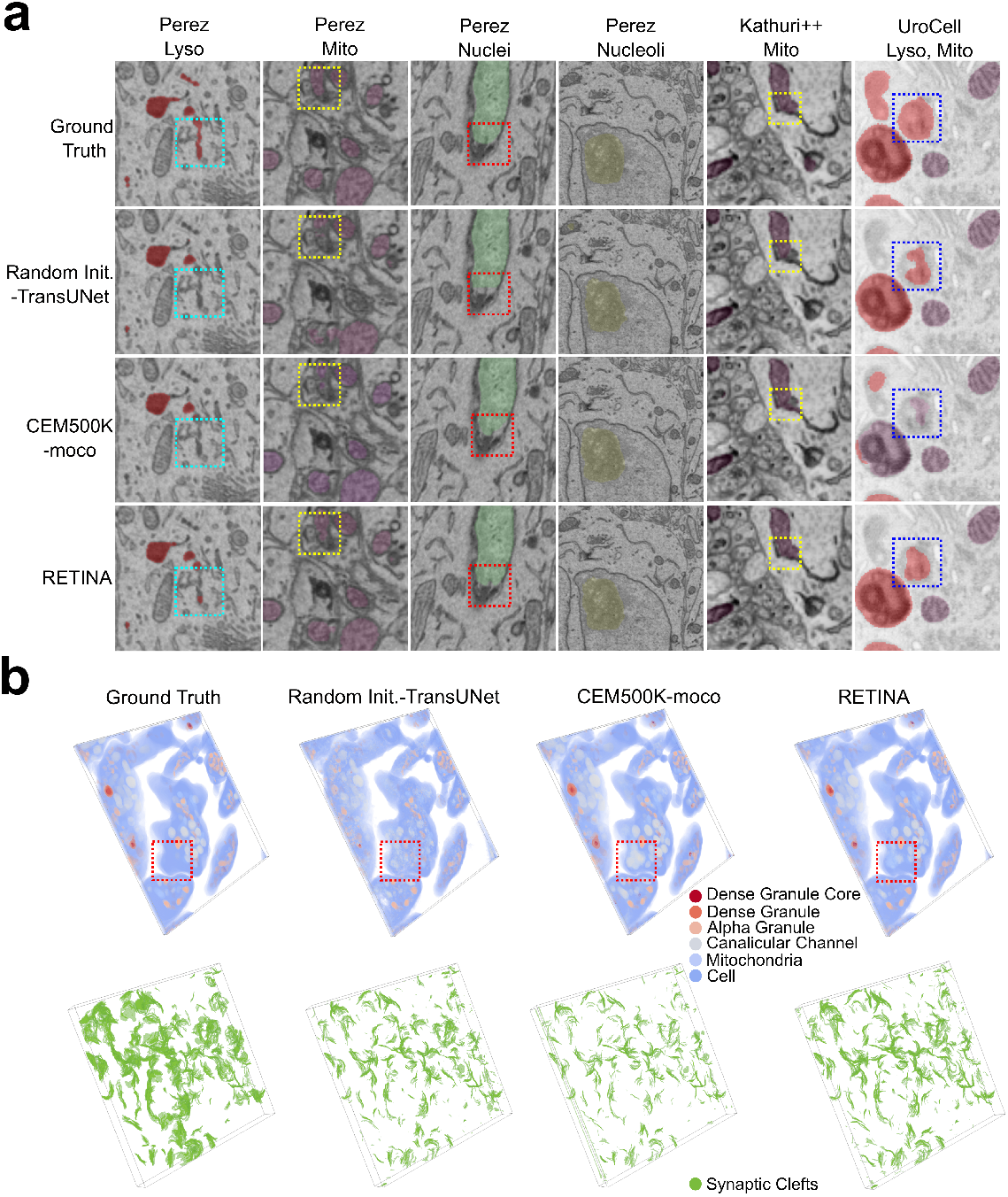
Visual comparison of segmentation. **a**, Example segmentations from all 2D benchmark datasets. The first row shows the ground truth; the second row represents the randomly initialized TransUNet; the third row depicts the MoCoV2 pre-trained model; the fourth row corresponds to RETINA. Lysosomes are shown in red, mitochondria in purple, nuclei in green, and nucleoli in yellow. **b**, Example segmentations from 3D benchmark datasets. Labels for the Guay and CREMI Synaptic Clefts datasets are shown in the first and second rows, respectively. The dashed-line squares highlight regions where RETINA demonstrates improved segmentation accuracy. Abbreviations: Lyso, Lysosomes; Mito, Mitochondria.

For 3D visualization, we reconstructed 3D label maps for the Guay and CREMI Synaptic Clefts datasets by stacking all inferred 2D slices along the z-axis. In the Guay benchmark, the regions highlighted with red squares reveal that both the TransUNet and MoCoV2 pre-trained models generate numerous false labels (Figure 4b). Specifically, the ground truth within these squares predominantly represents a pure cell region (colored in blue) without additional labeled cellular material. However, these two models mistakenly assign substantial canalicular channels (colored in grey) and some mitochondria (colored in light blue) to this region (Figure 4b). In contrast, RETINA’s result (last column in Figure 4b) exhibits fewer misassignments, with more of the region correctly classified as the pure cell category (colored in blue). For the challenging CREMI Synaptic Clefts dataset (second row in Figure 4b), where benchmark models have underperformed (Conrad and Narayan, 2021) (Table 1). RETINA’s inference reveals more and more accurate synaptic cleft labels (colored in green) compared to the other models. Additionally, the randomly initialized TransUNet (second column) identifies more synaptic clefts than the MoCoV2 pre-trained model (third column) suggesting that the improvement in RETINA is largely attributable to the TransUNet architecture, which enhances the capture of detailed synaptic cleft features. In conclusion, both 2D and 3D segmentation visualizations demonstrate that RETINA effectively captures organelle features, correcting mismatches in multi-class segmentation tasks.

### 3.2 RETINA performs well benefiting from the CEM500K dataset

One of the benefits of RETINA is its ability to pre-learn features from the CEM500K dataset. To further explore the importance of pre-training on CEM500K, we experimented with different pre-training dataset selections, including ImageNet, CEM500K, and a combination of both (Table S5), while keeping other parameters constant. Given that ImageNet is a labeled dataset, pre-training was supervised and applied to the TransUNet model. For the model pre-trained on both ImageNet and CEM500K, it was first pre-trained on ImageNet, and then the parameters were transferred to the reconstruction-based architecture (Figure 1a) for further pre-training on the CEM500K dataset. Pre-training on the CEM500K dataset results in better performance, as indicated by higher IoU scores on the CREMI Synaptic Clefts, Guay, Kathuri++, and Perez datasets, whereas ImageNet pre-training is superior for the UroCell dataset (Table S5). Our findings show that CEM500K pre-training generally outperforms ImageNet pre-training, with the exception of the UroCell dataset, where ImageNet pre-training achieves superior results. The diminished performance of the CEM500K pre-trained model on UroCell suggests that the domain-specific advantages of CEM500K are less applicable to this dataset compared with the other benchmark datasets. Additionally, a combined pre-training approach using both ImageNet and CEM500K does not surpass CEM500K pre-training alone across the benchmark datasets (Table S5), with only a slight improvement found for mitochondrial segmentation in the Perez benchmark. Thus, combined pre-training does not offer substantial benefits, CEM500K pre-training remains a robust strategy for enhancing segmentation accuracy across most datasets.

## 4 DISCUSSION

RETINA enhances the accuracy of segmentation on EM-related cellular images, benefiting from two key aspects. First, RETINA uses a TransUNet backbone, with its hybrid encoder (comprising convolutional and transformer layers) achieving both local and global image feature extraction. Second, it is pre-trained on the CEM500K dataset, a diverse and non-redundant EM image dataset, that helps RETINA to pre-learn EM-related features and infer the nanoscale objects accurately after fine-tuning on the benchmarks.

Given that CEM500K is an unlabeled dataset, an unsupervised pre-training design is required. Initially, instead of the reconstruction-based unsupervised pre-training method applied for RETINA, we employed SimCLR due to its proven effectiveness in pre-training convolutional layers (Chen et al., 2020a). After pre-training, we transferred the pre-trained convolutional layers within the encoder to the fine-tuning phase. However, this approach did not yield any improvement over the randomly initialized TransUNet. Similarly, using masked autoencoders (He et al., 2022) to pre-train only the transformer layers resulted in poor outcomes. These findings suggest that pre-training only parts of the hybrid encoder is ineffective; instead, the entire encoder should be pre-trained. The complexity of the hybrid encoder makes it challenging to achieve good segmentation performance on EM images with pre-training, as the hybrid structure requires integration, consistency, and transition within the encoder (Li et al., 2024). In response, we developed a reconstruction-based pre-training method that recovers augmented images to their original state, involving both convolutional and transformer layers, thereby enabling consistent pre-training of the hybrid model. This reconstruction-based method should be generalizable to other hybrid models, such as U-Mamba (Ma et al., 2024a), for pre-training on unlabeled datasets.

The similar model structure between the RETINA pre-training and fine-tuning phases should enable seamless transferring of the pre-trained layers from the corresponding pre-training phase to the fine-tuning phase. However, transferring all parameters did not result in improved performance; instead, the best results were achieved by transferring only the transformer layer parameters. This may be because more adjustable parameters are required to accommodate the information learned during pre-training and fine-tuning. In the RETINA design, the convolutional layers remain unfrozen during fine-tuning, allowing them to adapt to the fine-tuning dataset, while only the transformer layers are frozen. This approach enables the convolutional layers to adjust and learn new features during fine-tuning, making RETINA more flexible and adaptable. At the same time, the features learned from CEM500K during pre-training are preserved by freezing the transformer layers, preventing them from being altered during fine-tuning. This strategy appears to balance the retention of pre-trained features with the adaptation to new data, resulting in more accurate segmentation.

Given that augmentation can increase the diversity of the dataset and modify the difficulty of the training task, it needs to be carefully designed (Cubuk et al., 2018). The set of augmentation operations can be customized based on different pre-training datasets and downstream tasks to maximize the benefits obtained from pre-training. For RETINA, we employed random flips, rotations, brightness, and contrast adjustments as standard methods to augment the dataset. Additionally, adding noise and blur increased the reconstruction difficulty. Inspired by the masked autoencoder model (He et al., 2022), we also randomly masked parts of the input images to enhance feature learning. Effective augmentation ensures that RETINA’s deeper layers are thoroughly trained. Given the skip connection design of TransUNet, insufficiently robust augmentation may cause the input image vectors to bypass the deep layers, leading to quick convergence without adequately updating the deeper layers (Figure 1a), resulting in insufficient pre-training and no MSE loss improvement.

The choice of pre-training dataset can significantly impact the ultimate accuracy of segmentation. Although the CEM500K dataset comprises 0.5 × 10^6^ diverse and informative cellular EM images, there could still be space to generate a more powerful pre-training dataset for different tasks. When the downstream task involves images similar to the pre-training images, the model can more efficiently capture relevant features, potentially outperforming models pre-trained on large but unrelated datasets (Krishna et al., 2022). The image similarity between the pre-training dataset and the fine-tuning dataset can be influenced not only by the type of target objects they share but also by their optical properties, such as brightness, contrast, and white balance (Fish, 2009). This may explain why pre-training on CEM500K is less effective for the UroCell dataset compared to supervised pre-training on large ImageNet datasets (Table S5), as UroCell images have higher brightness compared to all other benchmarks (Figures 2b and 4a). Therefore, there is potential to create more suitable pre-training datasets that balance generalization and task-specific features based on downstream tasks. Nevertheless, RETINA’s hybrid encoder structure helps mitigate the limitations of pre-training datasets. For instance, in the CREMI Synaptic Clefts benchmark, the MoCoV2 pre-trained UNet-ResNet50 model improved the IoU score from 0.000 to 0.246 (Table 1), while RETINA, without any pre-training on CEM500K, achieved an even higher IoU score of 0.294. This suggests that the hybrid encoder structure can be more important than pre-training, especially when the features of the pre-training dataset are not well-aligned with those of the fine-tuning dataset, and the benefits of pre-training may be diminished in such cases.

One important benefit of pre-training is that it can reduce the time required for training downstream tasks (Minaee et al., 2021) (see timing information in Supplementary Material). In our experiments, we observe that RETINA converges faster compared to the benchmark models without pre-training (Figure 3a). In our CREMI Synaptic Clefts, Perez, and UroCell benchmarks, RETINA demonstrates rapid convergence within just a few hundred steps. The required number of fine-tuning iterations may vary depending on the complexity of the tasks and the relevance of features between the pre-training and fine-tuning datasets. In our experimental design, we set the number of fine-tuning iterations based on prior work (Conrad and Narayan, 2021) to ensure a consistent basis for performance comparisons. For future applications of RETINA, fewer fine-tuning steps could be used to reduce runtime, making the model more efficient.

As EM datasets grow, it is likely that more powerful pre-training datasets can be developed for various specific tasks to achieve efficient pre-training. Additionally, to achieve more general performance, pre-training datasets are typically very large, making manual mask curation difficult. Therefore, effective pre-training methods that do not require labels, as we propose here, will need to be developed. Although our reconstruction-based model shows good performance with the TransUNet backbone, other hybrid encoder combinations, such as U-Mamba (Ma et al., 2024a) and SwinUNETR (Hatamizadeh et al., 2021), warrant exploration to further improve performance. Hopefully, continued research in this area will lead to automated segmentation reaching the accuracy of expert manual segmentation.

## Supporting information

RETINA supplementary material

## AUTHOR CONTRIBUTIONS

**Cheng Xing**: Conceptualization, Methodology, Validation, Formal analysis, Investigation, Data Curation, Writing - Original Draft, Writing - Review & Editing, Visualization. **Ronald Xie**: Conceptualization, Supervision. **Gary D. Bader**: Resources, Writing - Review & Editing, Supervision, Project administration, Funding acquisition.

## ACKNOWLEDGMENTS

All authors thank the Digital Research Alliance of Canada for high-performance computing access.

## SUPPLEMENTAL DATA

Supplementary material has been prepared.

## DATA AVAILABILITY

The benchmark datasets and the pre-trained weights are available at Zenodo.

## CODE AVAILABILITY

The full code of RETINA implementation is available at GitHub.

