## Supplementary material for "RETINA: Reconstruction-based Pre-Trained Enhanced TransUNet for Electron Microscopy Segmentation on the CEM500K Dataset": RETINA supplementary material

#### 1 IMPLEMENTATION

##### 1.1 RETINA Pre-training Methods

RETINA preprocesses each input image with augmentation. Each input is duplicated once to form a pair: one is designated as ImageA and the other as ImageB. ImageB is saved as the original image and damaging augmentations are applied to ImageA. The augmentations are classified into two sets, implemented using the 'albumentations' Python package (Buslaev et al., 2020). The first set comprises 'HorizontalFlip', 'VerticalFlip', 'Rotate', and 'RandomResizedCrop', applied to both images in the pair. These don't damage any pixels. The second set includes 'RandomBrightnessContrast', 'GaussNoise', 'GaussianBlur', and 'CoarseDropout'. These damage pixels, so are applied only to ImageA, with ImageB serving as the reference for comparison. The parameters for each augmentation function are detailed in Table S1.

RETINA encodes ImageA using a combination of convolutional and Transformer layers, following the TransUNet architecture. Among the various TransUNet configurations, we selected the ResNet50 and ViT-B/16 configuration because ViT-B/16, when combined with ResNet50, has demonstrated excellent performance in previous studies (Chen et al., 2021). Additionally, ViT-B/16 offers a more compact structure compared to the L/16 variant, and we further optimized the model by reducing the number of parameters (Table S2) to balance high accuracy with a reduced training burden. The Transformer encoder consists of Multihead Self-Attention (MSA) layers and Multi-Layer Perceptron (MLP) layers (Dosovitskiy et al., 2020), with the output after the encoder given by:

$$z_\ell = \mathrm{MLP}(\mathrm{LN}(\mathrm{MSA}(\mathrm{LN}(z_{\ell-1}))) + z_{\ell-1}) + \mathrm{MSA}(\mathrm{LN}(z_{\ell-1})) + z_{\ell-1}, \quad \text{(S1)}$$

where LN() denotes the normalization operator, and $z_\ell$ is the encoded representation. The architecture is illustrated in Figure 1a. Additionally, skip-connections are set to 3 to enable efficient feature aggregation at different resolution levels. After decoding the features obtained from the encoder, a 3x3 convolutional layer is applied to recover the decoded feature instead of the segmentation head used in the original TransUNet architecture. Finally, the MSE loss is calculated between the reconstructed output and ImageB. The overall parameter settings are detailed in Table S2. RETINA is implemented with PyTorch in a Python 3.10.2 environment and pre-trained for 200 epochs on Nvidia A100 GPUs on the Digital Research Alliance of Canada's Narval cluster. The training was distributed across 4 nodes, each with 4 GPUs, and completed in less than 2 days.

##### 1.2 RETINA Fine-tuning Methods

The volume images were split into 2D images as input for the model. To transfer the pre-trained parameters, we maintain the same encoder and decoder parameter settings during fine-tuning as in the pre-training phase (Table S3). The pre-trained transformer parameters were transferred to the fine-tuning transformer layers and frozen during training. After decoding the features from the latent space, they were processed by a segmentation head composed of a convolutional layer and a bilinear upsampling layer. For multi-class segmentation tasks, the final loss minimized by RETINA is given by (Chen et al., 2021):

$$L = \lambda L_{\mathrm{CE}} + (1-\lambda) L_{\mathrm{D}} \tag{S2}$$

where $\lambda$ is in the range [0,1], $L_{\mathrm{CE}}$ stands for cross-entropy loss, and $L_{\mathrm{D}}$ represents dice loss. For binary tasks, the loss is given by (Chen et al., 2021):

$$L = \lambda L_{\mathrm{BCEL}} + (1-\lambda) L_{\mathrm{BD}} \tag{S3}$$

$$L_{\mathrm{BCEL}} = -\frac{1}{N}\sum_{i=1}^{N}\left[y_i \log(p_i) + (1-y_i)\log(1-p_i)\right] \tag{S4}$$

where $L_{\mathrm{BD}}$ is the binary dice loss, N is the number of pixels, $y_i$ is the true label for the $i^{th}$ pixel, and $p_i$ is the predicted probability that the pixel belongs to the positive class. In both multi-class and binary tasks, $\lambda$ is set to 0.5. Fine-tuning was implemented on Nvidia T4 GPUs on the Digital Research Alliance of Canada's Graham cluster, with multi-GPU computation deployed to reduce computation time.

##### 1.3 Inference Methods

All benchmarks were inferred as 2D segmentation tasks, slice by slice. For UroCell, predictions were made on the xy, yz, and xz cross-sections due to its isotropic voxels, and the scores from each cross-section were averaged to obtain the final result. The remaining benchmarks were predicted along a single direction Evaluation followed the CEM500K publication guidelines (Conrad and Narayan, 2021). For the Guay dataset, the first volume was used for training, and the second for testing, excluding the third volume due to partial labeling. Evaluation on all benchmark data is based on the Intersection over Union (IoU) metric, defined as:

$$IoU = \frac{TP}{TP + FN + FP} \tag{S5}$$

where $TP$ is the number of true positive predicted pixels, $FN$ is the number of false negative predicted pixels, and $FP$ is the number of false positive predicted pixels. All benchmark results were reimplemented using the same parameters as the CEM500K (Conrad and Narayan, 2021) publication. RETINA uses the sliding window method for inference (Cardoso et al., 2022), with adjacent regions of prediction partially overlapped to reduce missegmentation at the borders. Inference was conducted on Nvidia T4 GPUs on the Graham cluster.

##### 1.4 Benchmark model implementation

We chose three models for benchmarking: TransUNet with randomly initialized parameters, UNet-ResNet50 with randomly initialized parameters, and UNet-ResNet50 pre-trained on CEM500K with MoCoV2. The TransUNet with randomly initialized parameters followed the same training and inference methods as described above. For the UNet-ResNet50 pre-trained on CEM500K with MoCoV2, we adhered to the details provided in (Conrad and Narayan, 2021). To maintain consistency, we applied their pre-trained model with 200 epochs by MoCoV2 on CEM500K and transferred the parameters to the fine-tuning model. During fine-tuning on the benchmark training sets, we froze the encoder part of the model. The configurations for each benchmark were adopted from the provided configuration files (https://github.com/volume-em/cem-dataset) (Conrad and Narayan, 2021). Checkpoints were saved at specific intervals of iterations (Figure 3a) for inference. Fine-tuning was implemented on Nvidia T4 GPUs on the Graham cluster. For inference, each benchmark could use one of two inference Python scripts: one for 2D and one for 3D. We used the script consistent with the benchmark task described in Table S4. Finally, after obtaining the predicted images, the IoU scores were calculated.

#### 2 SUPPLEMENTARY TABLES

**Table S1.** RETINA augmentation function settings. The first column lists the function name. The second column shows the probability of each function being applied to the image. The third column indicates the augmentation set number to which each function belongs. Set 1 comprises the augmentation operations applied to both input images, ImageA and ImageB, as outlined in the RETINA pre-training methods section. Set 2 includes the operations applied exclusively to ImageA. The last column details the specific parameter settings for each function.

| Augmentation | Probability | Set number | Parameters |
|---|---|---|---|
| HorizontalFlip | 0.5 | 1 | None |
| VeriticalFlip | 0.5 | 1 | None |
| Rotate | 0.5 | 1 | limit=90 |
| RandomResizedCrop | 1.0 | 1 | height=width=224<br>scale=(0.08, 1),<br>ratio=(0.5, 1.5) |
| RandomBrightnessContrast | 1.0 | 2 | brightness_limit=0.3<br>contrast_limit=0.3 |
| GaussNoise | 1.0 | 2 | var_limit=(400, 1200) |
| GaussianBlur | 1.0 | 2 | Default |
| CoarseDropout | 1.0 | 2 | max_holes=32<br>max_height=16<br>max_width=16<br>min_holes=1<br>fill_value=0<br>always_apply=False |

**Table S2.** RETINA implementation parameters for pre-training.

| Config | Parameters |
|---|---|
| ResNet number of layers | (3, 4, 9) |
| ResNet width factor | 1 |
| ViT name | R50-VIT-B_16 |
| patch size | (16, 16) |
| hidden size | 192 |
| MLP dimension | 768 |
| number of heads | 12 |
| number of layers | 12 |
| attention dropout rate | 0.0 |
| transformer dropout rate | 0.1 |
| decoder channels | (256, 128, 64, 16) |
| skip channels | (512, 256, 64, 16) |
| number of skip | 3 |
| optimizer | Stochastic Gradient Descent |
| momentum | 0.9 |
| batch size | 256 |
| learning rate | 0.003 |
| weight decay | 0.0001 |

**Table S3.** RETINA implementation parameters for fine-tuning.

| Config | Parameters |
|---|---|
| ResNet number of layers | (3, 4, 9) |
| ResNet width factor | 1 |
| ViT name | R50-VIT-B_16 |
| patch size | (16, 16) |
| hidden size | 192 |
| MLP dimension | 768 |
| number of heads | 12 |
| number of layers | 12 |
| attention dropout rate | 0.0 |
| transformer dropout rate | 0.1 |
| decoder channels | (256, 128, 64, 16) |
| skip channels | (512, 256, 64, 16) |
| number of skip | 3 |
| optimizer | Stochastic gradient descent |
| momentum | 0.9 |
| batch size | 16 |
| learning rate | 0.003 |
| weight decay | 0.1 |
| learning rate policy | Polynomial decay |

**Table S4.** Characteristics of the benchmark datasets. Abbreviations: ssTEM, serial section transmission electron microscopy; SBFSEM, serial block-face scanning electron microscopy; AT-SEM, array tomography scanning electron microscopy; FIBSEM, focused ion beam scanning electron microscopy.

| Benchmark | Microscopy type | Training set # of images | Training set image size | Testing set # of images | Testing set image size | Segmentation class(es) |
|---|---|---|---|---|---|---|
| CREMI Synaptic Clefts | ssTEM | 248 | 1250x1250 | 125 | 1250x1250 | Synaptic Clefts |
| Guay | SBFSEM | 49 | 800x800 | 23 | 800x800 | Mitochondria<br>Canalicular channels<br>Alpha granules<br>Dense granules<br>Dense granule cores |
| Kasthuri++ | AT-SEM | 85 | 1463x1613 | 75 | 1334x1553 | Mitochondria |
| Perez | SBEM | 50 | 500x500 | 40 | 1000x1000 | Mitochondria<br>Lysosomes<br>Nuclei<br>Nucleoli |
| UroCell | FIBSEM | 3006 | 202x256 | 753 | 244x256 | Mitochondria<br>Lysosomes |

**Table S5.** Comparison of IoU scores for models pre-trained on different datasets. The models include randomly initialized TransUNet, TransUNet pre-trained on ImageNet, RETINA pre-trained on CEM500K, and RETINA using ImageNet pre-trained parameters followed by pre-training on CEM500K. These models were fine-tuned and evaluated on all benchmark datasets listed in the first column. The IoU scores represent the best performance achieved with the specific number of training iterations shown in the second column. Lysosomes, mitochondria, nuclei, and nucleoli within Perez benchmark are listed separately. Mean values of three independent runs are reported.

| **Benchmark** | **Training Iterations** | **Random Init.** | **ImageNet** | **CEM500K** | **ImageNet & CEM500K** |
|---|---|---|---|---|---|
| CREMI S.C. | 5000 | 0.294 | 0.314 | **0.327** | 0.305 |
| Guay | 1000 | 0.223 | 0.390 | **0.513** | 0.359 |
| Kasthuri++ | 10000 | 0.894 | 0.910 | **0.916** | 0.914 |
| UroCell | 1000 | 0.579 | **0.649** | 0.610 | 0.596 |
| Perez | 2500 | 0.893 | 0.909 | **0.919** | 0.914 |
| Lysosomes | – | 0.845 | 0.844 | **0.885** | 0.862 |
| Mitochondria | – | 0.847 | 0.889 | 0.890 | **0.892** |
| Nuclei | – | 0.990 | 0.990 | **0.991** | 0.991 |
| Nucleoli | – | 0.891 | **0.914** | 0.910 | 0.910 |

### 3 SUPPLEMENTARY FIGURES

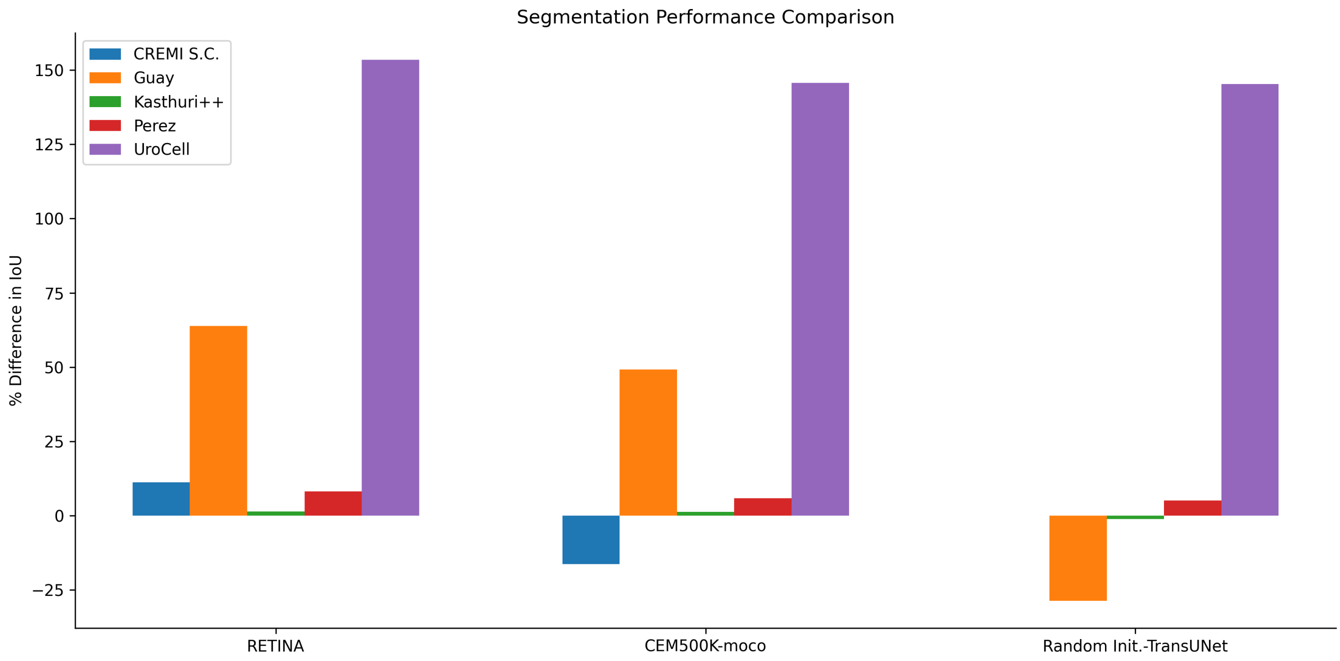


**Figure S1.** Percent difference in IoU (Intersection over Union) between RETINA and other models in bar plot. Randomly initialized UNet-ResNet50 serves as the baseline for all benchmark datasets except for CREMI S.C. (Synaptic Clefts), where the randomly initialized TransUNet model is the baseline.
